# Differential evolutionary and ecological patterns in eye loss between parallel visual systems in spiders

**DOI:** 10.64898/2026.05.08.723754

**Authors:** M. Antonio Galan-Sanchez, F. Andres Rivera-Quiroz, Lauren Sumner-Rooney

## Abstract

Eye loss has long fascinated evolutionary biologists and occurs across the animal kingdom. Spiders have two parallel visual systems — two primary and six secondary eyes — but eye losses, leaving six, four, two, or no eyes, have occurred in multiple lineages. Despite their significance, reports of eye loss are scattered, limiting broader analysis. Here we present the first comprehensive analysis of eye loss across all known spider lineages. We show that eye loss occurs in ∼12% of extant species, mainly within the clade Synspermiata. Six-eyed spiders are most common (>5,300 species), while four-eyed, two-eyed, and eyeless forms are rarer and often linked to troglobitic lifestyles. Principal eye loss is widespread, occurring in 49 families across nearly all major lineages. Using a recent phylogeny of the order Araneae, we demonstrate a strong correlation between eye loss and occupancy of low-light environments, but this is complicated by differential effects across eye types and phylogenetic groups through geological time. These findings reveal striking lability in eye number and lay groundwork for future research into ecological, developmental, and neurological drivers of eye loss. [hidden Markov models, ancestral state reconstruction, Araneae, discrete character evolution, principal eyes, secondary eyes, low light environments].

---

The evolution of eye loss is a recurrent theme in evolutionary ecology, occurring across dark habitats including caves, burrows, and the deep sea. The underlying mechanisms and evolutionary dynamics of eye loss have been studied in systems including crustaceans (Protas et al. 2011; Syme and Oakley 2012; Re et al. 2018), gastropods (Sumner-Rooney et al. 2016; Williams et al. 2022), bivalves (Malkowsky and Götze 2014; Audino et al. 2020, 2022), mammals (Nikitina et al. 2004), annelids (Gonzalez et al. 2018), and fishes (Wilkens and Strecker 2003; Jeffery 2009; Soares and Niemiller 2013). As most cases are limited to a single pair of eyes, eye loss is total, but animals with larger visual systems may not lose all eye simultaneously, potentially offering more detailed insights to the evolutionary causes and correlates of eye loss.

Spiders have some of the most diverse and sophisticated visual systems among arthropods. Most spiders have eight eyes, of two morphologically and developmentally distinct types: one pair of principal eyes and three pairs of secondary eyes, which are likely homologous to the ocelli and compound eyes, respectively, of extant hexapods (Paulus 1979; Samadi et al. 2015; Schomburg et al. 2015; Miether and Dunlop 2016). These parallel visual systems have been a feature of the arthropod body plan for over 500 Ma (Paulus 1979; Foelix 2011; Samadi et al. 2015; Miether and Dunlop 2016). In spiders each pair of eyes is named for its position on the carapace: the anterior median (AME; the principal eyes), anterior lateral (ALE), posterior median (PME), and posterior lateral (PLE) eyes (all secondary eyes) (Foelix 2011; Samadi et al. 2015; Schomburg et al. 2015). Spiders exhibit astonishing diversity in eye size, shape, position, and orientation (Morehouse et al. 2017; Morehouse 2020), and the modularity afforded by two types and four pairs of eyes has yielded an array of visual capabilities across both taxa and eye pairs; fields of view (Land 1985), spatial and temporal resolution, and contrast (Land and Barth 1992; Clemente et al. 2005, 2010; Insausti et al. 2012), polarization (Kovoor et al. 1993; Dacke et al. 2001), and wavelength sensitivities (Hu et al. 2012; Zurek et al. 2015) vary widely and can be uniquely tuned to behavioural and ecological needs (Morehouse 2020; Winsor et al. 2023).

Despite diversification into virtually every terrestrial environment, the eight-eyed blueprint has remained unchanged over nearly 400 million years (Selden et al. 2008; Dunlop 2022). This is in striking contrast to the variable number of lateral eyes found in other arachnids (Munoz-Cuevas 1984, Miether and Dunlop 2016), particularly scorpions (Yang et al. 2013; Loria and Prendini 2014), and in myriapods (Munoz-Cuevas 1984; Sombke and Müller 2023). However, eye losses have been described at varying phylogenetic depths in spiders, including species with six, four, two, and no eyes. Most records are scattered across the taxonomic literature and reporting of eye number is patchy, with some exceptions within specific families (Gertsch 1992; Cokendolpher 2004; Paquin and Dupérré 2009; Lin and Li 2010; Hedin et al. 2018; Huber 2018; Aharon et al. 2023; Wang et al. 2023). Together, the modularity and the conserved blueprint of spider visual systems offer an exceptional model to understand the evolutionary dynamics, ecological correlates, and underlying molecular and developmental mechanisms of eye loss. The loss of different types, pairs, or numbers of eyes may provide additional resolution in examining the evolution of loss in different ecological and phylogenetic contexts. Identifying the types of eye losses, where they occur in spider phylogeny, and potential links to ecological pressures, is essential to allocating efforts to understand the evolution of eye loss.

To address this need, we assembled a comprehensive dataset of eye number for the entire order Araneae. We extracted morphological, geographical, and microhabitat data for 7,620 species from >2,000 publications, capturing eye number and substantial reductions in size for both males and females. To reconstruct the evolutionary history of eye loss and its relation to ecology, we implemented models of discrete trait evolution and tested their correlation to light habitats. Our results show that eye loss is extremely common, reversible, and associated with low-light environments, but that its prevalence and evolutionary age vary across eye types and phylogenetic groups.

## MATERIAL AND METHODS

### Database assembly

We performed a comprehensive manual search of eye modifications across the taxonomic literature representing the whole Order Araneae. This search was not exhaustive at the species level but aimed to cover the largest number of families possible. Our search started by documenting the eye morphological variation at a family level, considering 134 of the 138 valid families according to the World Spider Catalog (WSC 2025) (Supplementary Dataset 1). We collected information about eye variation from family-level keys, descriptions and other specialized literature, as well as conducting Google Scholar searches with the family name + eye keywords (e.g. Tetrablemmidae “four eyes”) for each family and potential number of eyes to find potential cases of isolated or more recent descriptions that were missed in the keys. After this, a deeper search at the species level was performed for every family where any type of eye loss had been recorded. Information about the number, relative size of the eyes, habitat, geographical distribution and potential sex differences was directly extracted or inferred from the original species descriptions in taxonomic literature. We reviewed more than 1,300 publications, generating a database that represented 7,620 species and 56 spider families (Supplementary Dataset 2).

Due to the heterogeneity, ambiguity, or unavailability in the habitat information, we classified the habitats in ten categories to cover the principal microhabitats that spiders occupy (Supplementary Data 1). Then, these categories were split in two main groups accounting for the type of environment, i.e., degree of light exposure they receive, that we later used to test for correlation between eye loss and these environments.

### Phylogenetic comparative methods

#### Models of evolution for eye loss and ancestral state reconstruction

We implemented four character-coding schemes to explore the evolution of eye loss in spiders (see Supplementary Data 1 for taxon selection and treatment). We first assessed the hypothesis of eye loss by examining the origin of the secondary and primary eye systems, focusing on transitions from the presence of both visual systems (PE+3SE) to the absence of one of them (i.e., 3SE or PE) or both altogether (no eyes). Second, we explored the origin of specific eye configurations by distinguishing losses in the principal and secondary systems and the combination of the retained eyes. We identified eight distinct configurations: four involving the presence of the principal eye system, namely, PE+3SE; PE+2SE; PE+1SE; PE, three including only the secondary eye system (3SE, 2SE and 1SE) and one representing the eyeless type (no eyes) (Fig. 5, Appendix 1). We also analysed the two eye types independently with the absence/presence for the PE and multistate for the SE (absent, 1, 2, or 3 pairs). Lastly, we evaluated the presence or absence of the four eye pairs independently (AME, ALE, PME, PLE).

Then, we used the eight configurations to hypothesise two possible scenarios for eye loss: (1) eye number is reduced by losing one pair at a time on each evolutionary event or (2) loss occurs through the disappearance of one or more eye pairs in a single evolutionary event (Fig. 5). Given the distinct developmental origin of the principal eye system and its frequent absence/presence within the same spider group, we further hypothesize that this eye pair may have been regained in some lineages (e.g., Synspermiata).

We used the corHMM v2.8 R package (Beaulieu et al. 2017) to fit and compare these different coding schemes and hypotheses by employing Standard Markov (Mk) and Hidden Markov Models (HMM) (Supplementary Table S2, Supplementary Code 1). The latter allows for variation in transition rates throughout the tree by implementing “hidden” states within the observed states (Tarasov 2019; Boyko & Beaulieu 2021). We fit customized models with no hidden states, and with one hidden state (1 and 2 rate categories, respectively) on the three coding schemes (eight- and four-character states schemes and the four eye pairs independently). Models were built by either placing restrictions on the transition matrix to prevent transitions not allowed in a given hypothesis (e.g., presence to absence only) or permitting these changes to occur (i.e., loss of AME and vice versa). Transition rates between pairs of states were either all estimated independently (all-rates-different model, ARD) or constrained to be the same (equal-rates model, ER). Furthermore, we fit hybrid hidden rate models where one rate class was a restricted model and the other was either an ER model or an ARD model, and others where the rate class was ER and the other ARD. A combination of the restricted, ER, and ARD was implemented on the three hidden states model. For the purpose of comparison, we also fit ER, SYM, and ARD models without any constriction in the transition matrix on the four-character states coding scheme (PE + 3SE, 3SE, PE, no eyes). Finally, each model was fitted with two different root priors— *“equal”* where all states have equal probabilities occurring at the root (Schluter et al. 1997), and *“fitzjohn”* which weights each root state according to its probability of giving rise to the extant data, given the model parameters and the tree (Fitzjohn et al. 2009). We fit a total of 66 models. In order to compare model performance and avoid overparameterization, for each model we calculated the corresponding Akaike information criterion (AIC) (Beaulieu et al 2013) and the weighted Akaike information criterion (AICc) values. We computed marginal ancestral state reconstructions on the best-fit Mk or HMM models (determined by weighted AIC of the maximum likelihood estimates) as implemented in the function *corHMM*. We considered an ancestral node to likely be in a given state if the posterior probability for presence was > 70%.

#### Transitions through time

The rate of each state’s transition through evolutionary time was calculated to test whether certain transitions were more common during specific periods in evolutionary history (Supplementary Code 1). The tree was first divided into 30 and 60 equal bins (the total edge length of the phylogenetic tree is 304.06 million years), resulting in time blocks of 10.1 and 5.06 million years each. The rate through time was calculated as the ratio of the mean number of changes in each time block and the total edge length of the tree after accounting for the number of lineages in that time block (Revell 2017). The rate through time was calculated using 1,000 stochastic maps for the best-fit transition models of three coding schemes (i.e., eye pairs, four states scheme and eight states scheme). R codes from Hughes et al. (2021) and Halali et al. (2025) were used/modified for this analysis.

#### Correlation tests

The *fitCorrelationtest* function of the corHMM package was implemented to test for correlation between eye pairs (e.g., AME vs. PME) and, between eye pairs (e.g., AME) or eye number (e.g., 0 eyes) and microhabitats (low-light environments) (Supplementary Code 1). The test fits multi-rate dependent and independent models on binary character datasets that employ standard and hidden Markov models, the latter allowing for heterogeneity of the analysed characters and the correlation between them (Boyko & Beaulieu 2023). For each model the Akaike information criterion (AIC) and the weighted Akaike information criterion (AICc) values were calculated.

#### Phylogenetic signal

We evaluated the degree of phylogenetic signal for the four eye pairs (i.e., AME, ALE, PME, PLE) estimating the D statistic using the *phylo.d* function in the caper v1.0.3 R package (Supplementary Code 1). The D value is the sum of state changes along branches for a binary character, and it varies from 0 to 1 (Fritz and Purvis 2010). Thus, small values (D < 0) indicate few state changes in sister clades, supporting a strong phylogenetic signal and a clumped pattern of a given character in the phylogeny, D = 0 indicates that a character evolves on the phylogeny following the Brownian model of evolution and D = 1 indicates that characters are over dispersed on the phylogenetic tree. We conducted 1,000 permutations on the calibrated tree to test whether the loss of a particular eye pair was significantly different from the Brownian model or a random pattern of evolution.

### Habitat analysis

In addition to the correlation tests, we also applied Mk models, and stochastic character mapping to further explore the evolution of eye number reduction and the ancestral microhabitats under a Bayesian approach (Supplementary Code 1). We discretised the absolute eye number (i.e., 8, 6, 4, 2, 0) and the microhabitat information into binary characters. Eye numbers were discretised in “eight eyes” vs. “six or fewer eyes” states and microhabitats were split out in “low-light environments” vs. “others” according to our habitat categorisation (Supplementary Data 1). We used the *fitMk* function of the phytools R package to fit standard Markov models on the two datasets implementing ER and ARD transition models. Each model was fitted with *“equal”* and *“fitzjohn”* root priors (see previous sections) with its corresponding Akaike information criterion (AIC) values. We then generated stochastic character maps on the best-fitting models using the *simmap* function (1,000 simulations). This mapping approach reconstructs the character histories by estimating changes along the branches in addition to calculating probabilities of states at nodes (Huelsenbeck et al. 2003). Finally, we used the *densityMap* function to visualise the posterior probabilities from the stochastic maps.

The ancestral state estimation for microhabitats was also analysed by fitting ER, SYM and ARD transition models on the *all microhabitat categories* dataset using the *fitMk* function. Stochastic character maps were generated on the best fitting model using the *simmap* function (1,000 simulations).

#### Sankey Charts

Subsets of the original dataset were visualized using the package ggalluvial (Brunson 2020) for ggplot2 in R (Supplementary Code 2). These subsets display the eye losses per clade and their proportion per habitat using the Microhabitat categorization mentioned above.

#### Heatmaps and Correspondence Analysis

Subsets of the original dataset were transformed into a contingency table and then visualized using the package pheatmap (Kolde 2025) in R (Supplementary Code 2). The colour scale was adjusted per column to highlight the highest number of occurrences per microhabitat for each clade (Appendix 4a), highest number of occurrences per eye number in relation to microhabitat (Fig. 4, Appendix 4b), and highest number of occurrences per clade in relation to the number of eyes (Appendix 4c). We used the same contingency tables with the *Correspondence Analysis* function in the FactoMineR (Lê and Husson 2008) package to explore relationships among our qualitative variables.

## RESULTS

### Eye loss across Araneae

Of >53,000 described, valid spider species across 135 families (WCS 2025), we found that at least 6,181 species have lost one or more eye pairs. This represents approximately 11% of species, belonging to 56 families distributed across the order (Fig. 1). These included spiders with six (5,389 species, 87% of cases), four (183 species, 3%), two (203 species, 3.3%), and no eyes (406 species, 6.6%) (Fig. 1). The AME (principal eyes) were lost in almost 90% of cases.

**Figure 1.**
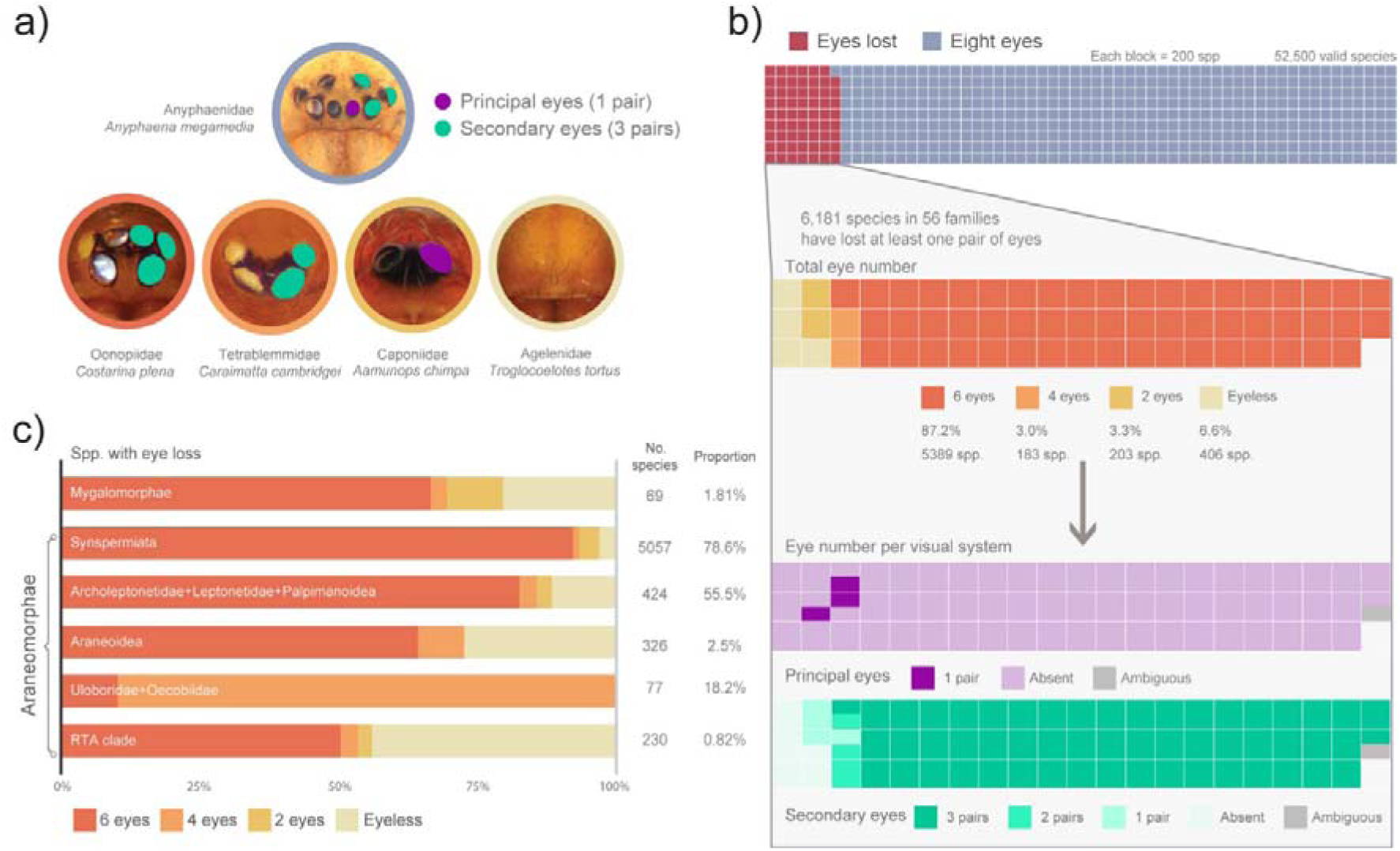
Diversity of eye number in spiders. a) Most spiders have four pairs of eyes split between two parallel visual systems: one pair of principal eyes (purple) and three pairs of secondary eyes (teal; top); however, variation in eye number appears in many families across spider phylogeny (bottom). b) Waffle plot showing the proportion of all described spider species exhibiting any form of eye loss (in red). Inset: Among species exhibiting eye loss, there is substantial variation in the total number of remaining eyes (i.e., 6–4–2–0; top) and in the number of remaining eyes by type (i.e. principal eyes present/absent and secondary eyes 1-3 pairs). c) Proportional representation of remaining eye numbers in species exhibiting loss, across major spider clades. Numbers to the right of each bar indicate the total number of species with eye loss and the percentage they represent within each clade.

The distribution of eye losses across spider phylogeny was broader than expected, affecting almost all major clades, from the plesiomorphic mygalomorphs to the derived RTA clade (Fig. 1c). However, losses were concentrated in certain clades, most notably in the early-diverging clade Synspermiata, where >5,000 of 6,200 described species (78.6%, across all 18 families) were affected, representing 81.8% of all reported spider species with eye loss. Other heavily affected groups included Palpimanoidea+(Leptonetidae+Archoleptonetidae) (424 species, 55.5%) and the family Uloboridae (77 species, 18.2%). Other major clades like Mygalomorphae, Araneoidea, and RTA had multiple records of eye losses, but these represented a tiny portion (<2.5%) of their respective species diversity.

We also found variation in the identity of the lost eyes. Although the principal eyes (PEs) were most commonly absent, some species lacked one or more pairs of secondary eyes (SEs) while the PEs persist (particularly in Caponiidae and Palpimanidae). This implies that there are two basic routes to eye loss, corresponding to the two eye types, and generates seven possible configurations: (1) PE+2SE; (2) PE+1SE; (3) PE only; (4) 3SE, (5) 2SE, (6) 1SE; (7) eyeless (Fig. 5, Appendix 1). A third route, resulting in rapid total eye loss, is considered separately.

Among six-eyed species (5,389 spp.), 99.1% lacked the AME (i.e., exhibited the 3SE state) and occurred in Synspermiata, Leptonetidae, and Archoleptonetidae. The PE+2SE configuration was only reported in 47 species across ten distantly related families, most notably Caponiidae (14 spp.) (Fig. 2, Appendix 1B.i). Generally, secondary eye loss affected the PMEs, except in Uloboridae, which apparently lose the ALEs.

**Figure 2.**
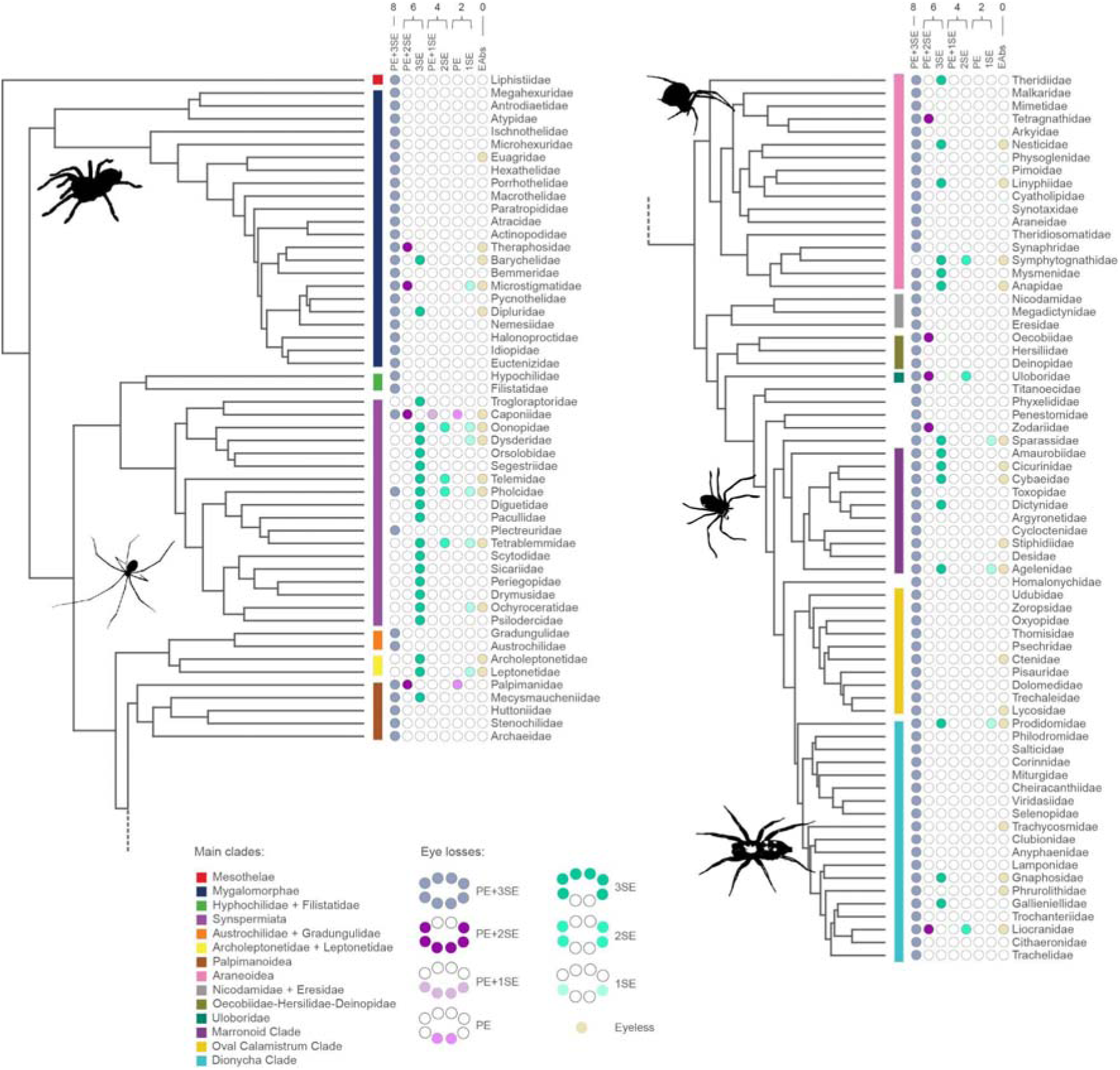
Family-level phylogeny derived from a 25% occupancy dataset of the ultraconserved elements (Kulkarni et al. 2023), showing the diversity of eye losses across the order. The cladogram depicts the eye losses in absolute number, -from eight to zero-, eye types (PE; principal eyes and SE; secondary eyes) and configurations (e.g., PE+3SE). Note that some families exhibit hypervariability in their visual systems. For instance, Caponiidae have five configurations, being one out of two families that preserves the PE and loses the SE. Other families with highly variable eyes are the Pholcidae (five configurations), and the Microstigmatidae, Oonopidae, Tetrablemmidae, Sparassidae, and Liocranidae (four configurations).

Four-eyed species (183 spp.) were rare. Most cases represented the 2SE configuration, particularly in Synspermiata (65 spp.; Fig. 2, Appendix 1E.i, ii, iii), as well as the families Uloboridae (69 spp.), Symphytognathidae (22 spp.), and Leptonetidae (19 spp.). The PE+1SE configuration was almost entirely restricted to Caponiidae (4 spp., Appendix 1D). One species in Dipluridae had a similar PE+1SE configuration, but the secondary eyes are curved and elongated, reminiscent of multiple, fused pairs (Appendix 2b).

Among two-eyed species (203 spp.), 61% retained the PEs and all such instances occurred in Caponiidae (121 spp., around 80% of caponiid species) and Palpimanidae (3 spp.) (Appendix 1F). The 1SE configuration appeared in five synspermiatan families, the families Leptonetidae and Microstigmatidae, and at least four families in the RTA clade (Appendix 1G).

The complete loss of eyes (406 spp.) occurred in 36 families across the order, representing 6.6% of species affected by eye loss, and 0.7% of all described spiders. The clades most affected were Synspermiata (153 spp.), the RTA clade (101 spp.), and Araneoidea (89 spp.).

### Evolution of eye loss

We reconstructed the evolutionary history of eye losses using models of discrete character evolution (summarized in Supplementary Table S1).

Our results support the typical eight-eyed arrangement (PE+3SE) being ancestral for spiders (pp: 0.99, 1-rate category *Reversal ER* model) (Fig. 3, Fig. 5, Appendix 1A). The best-scored models supported seven independent losses of the AME: one each at the base of Synspermiata (pp: 0.95), at the node Archoleptonetidae+Leptonetidae (pp: 0.96), within the family Mecysmaucheniidae (pp: 0.93) (within Palpimanoidea), in the family Symphytognathidae (pp: 0.99), and within the marronoid clade (pp: 0.99), and two losses within the family Anapidae (pp: 0.98, 0.99) (Fig. 3a, Fig. 5, Appendix 3). These reconstructions strongly supported the regain of the AME, i.e. reversal to eight eyes, three times independently in Synspermiata: at the bases of Caponiidae (pp: 0.99), and Plectreuridae (pp: 0.94), and within Pholcidae (pp: 0.97) (Fig. 3c, Appendix 1). For all secondary eye pairs, losses have evolved recently, appearing only at the tips of the phylogeny except the loss of the PMEs in the orb-weaver *Anapistula* (Fig. 3a, Appendix 3). Analysing the eight possible eye configurations as alternative states of a single character yielded the same patterns (Appendix 3), i.e. seven origins for the 3SE state, and 2SE as ancestral for *Anapistula*.

**Figure 3.**
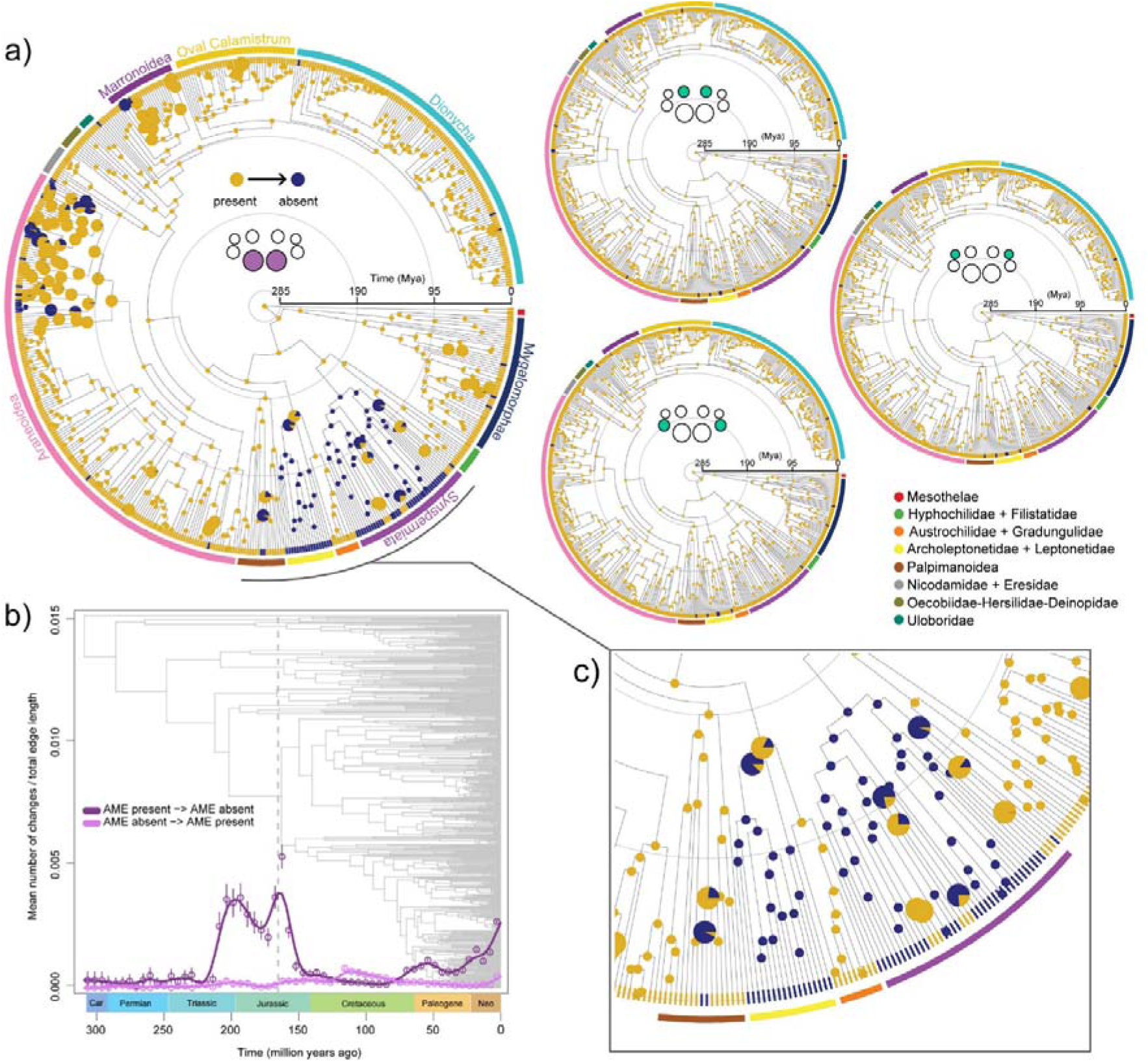
Maximum-likelihood phylogeny of spiders derived from a 25% occupancy dataset of the ultraconserved elements (Kulkarni et al. 2023), showing the ancestral state reconstruction of the four eye pairs and their transition through time. a) Marginal ancestral state estimation of the Anterior Median Eyes (AME) and the secondary eye pairs, PME (top), PLE (middle), and ALE (bottom). The AME loss has occurred seven times independently in different groups, particularly in Synspermiata and Archoleptonetidae + Leptonetidae, the former with a regain of this eye pair in three families: Caponiidae, Pholcidae and Plectreuridae (inset c). Smaller charts at nodes indicate probabilities > 90% for a given state. b) Transitions through time for the AME using stochastic maps. Each point (95% CI) represents the average number of transitions in a 5.06-million-year time block (see Methods), and the smoothed line represents the estimate from these points. Loss and gain of the AME are represented with dark and light purple, respectively. The vertical dotted line represents the diversification of the Synspermiata and the Archoleptonetidae + Leptonetidae clades, which occurred approximately ∼160 Ma. Transitions for PMEs, ALEs and PLEs are not shown because calculations for transitions resulted in zero values in multiple time blocks.

**Figure 4.**
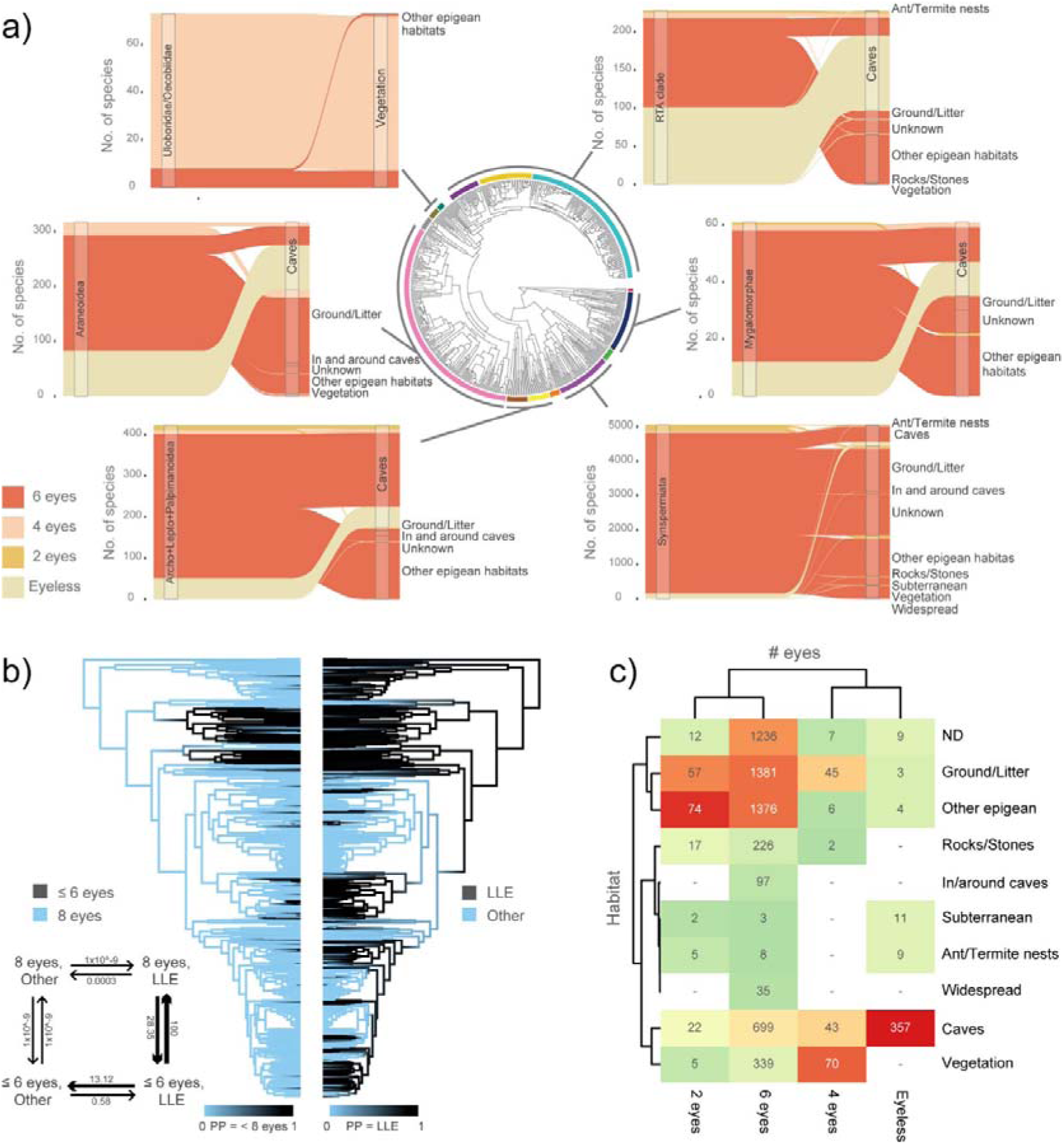
Eye number diversity and its correlation with habitat. a) Sankey charts showing the proportions of eye losses per habitat, per clade. b) Density maps of normal vs eye-reduced species plotted against low-light environments (LLE) vs other habitats. The arrow diagram depicts the transition rates between eye losses and the occupancy of LLEs under the Hidden-Correlated model (AIC: 724.68, AICc: 725.99). c) Heatmap showing eye number vs. the habitat, numbers in the cells indicate species. Note that virtually all eyeless species occur in caves regardless of phylogenetic affinity. Four eyes and two eyes are also mostly linked to caves and LLEs with two notable exceptions, the two eyes in Caponiidae (Synspermiata), and the four eyes in Uloboridae. Finally, the six-eyed state is very widespread and does not seem to be primarily driven by habitat.

**Figure 5.**
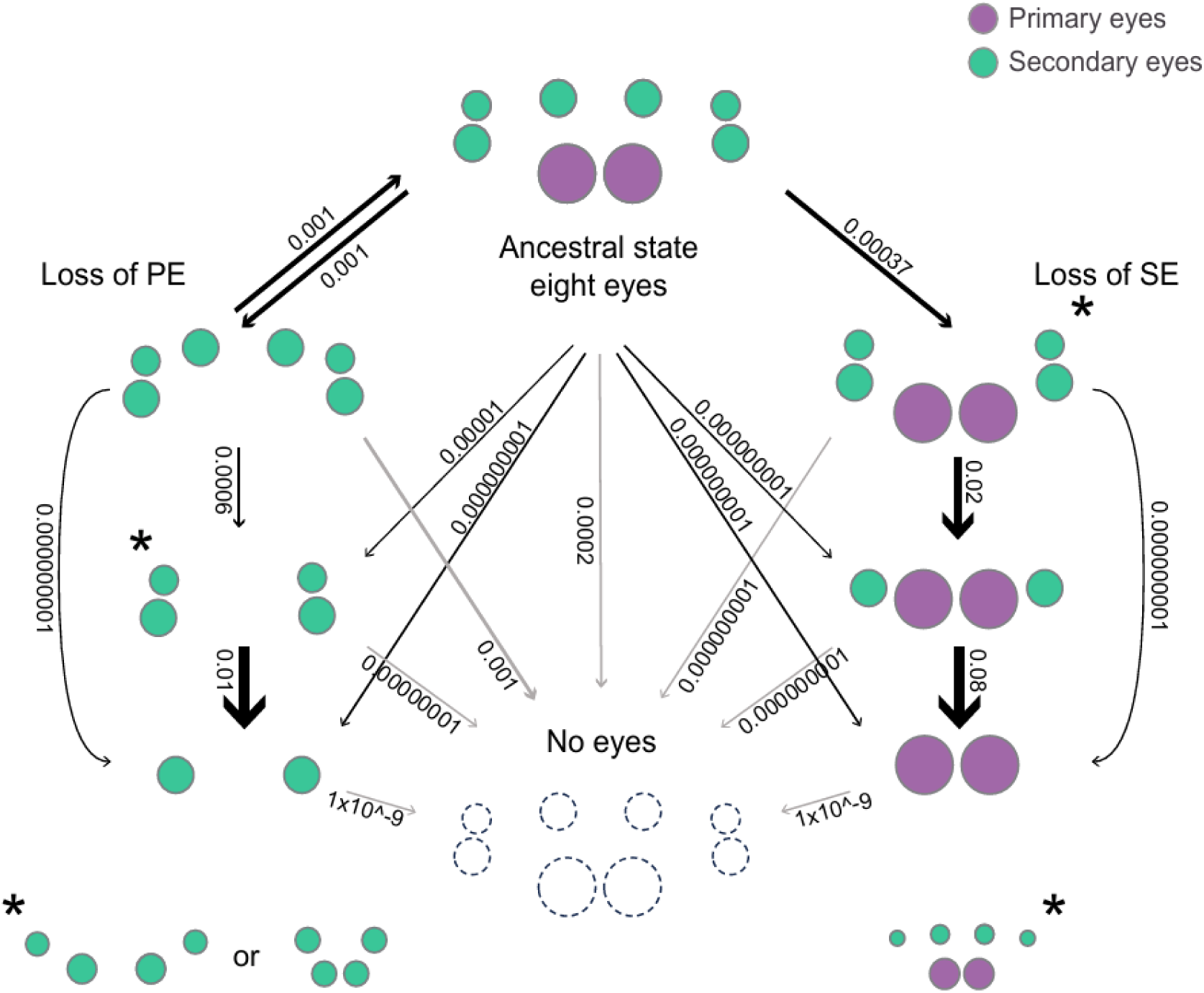
Hypothesis of the evolution of eye loss in spiders starting from the eight-eyes stereotypical configuration. In a first pathway the AME are lost (PE, left side of the schematic), resulting in a configuration composed solely by the secondary eye system (3SE). In the second pathway, the AME are retained while one pair of secondary eyes, most commonly the PMEs, is lost, producing a PE+2SE configuration (right side of the schematic). From these two major states, additional reductions may occur in either an ordered or unordered manner. In the 3SE condition spiders may lose one or two pairs yielding 2SE or 1SE configurations. Likewise, in the PE+2SE configuration, spiders lose one or two secondary eye pairs resulting in a PE+1SE or PE phenotype. A third pathway of eye loss involves the simultaneous loss of all eye pairs in a given configuration (grey arrows). Numbers in the arrows indicate the estimated transition rates under the 1 rate category *Reversal ER* model of the eight-states coding scheme (see methods). Asterisks show alternative arrangements of the 2SE and PE+2SE configurations. Abbreviations: PE; principal eyes, SE; secondary eyes.

We detected a clumped distribution of absent AME (*EstD*: -0.16), implying that they are more likely to be lost in closely related taxa. In contrast, secondary eye losses were randomly distributed across the phylogeny (*EstD* ALE: 0.94, *EstD* PLE: 0.86, *EstD* PME: 0.86). There was no correlation between losses of the AME and the secondary eyes, nor between the two lateral eye pairs (ALEs and PLEs), but we found support for a correlation between the losses of the PMEs and the lateral eye pairs (Supplementary Table S2).

Finally, we obtained transition rates between the presence and loss of individual eye pairs, and among eye configurations. The calculated rates of loss and gain of the AME varied from 0.001 (*Reversal SYM*) to 0.004 (*R2 Reversal ARD - R3 Reversal SYM*) and different rates (loss: 0.52, gain: 1.41) under the *R1 No reversal - R2 Reversal ARD* model, which were equally supported by AIC and AICc (Supplementary Table S1). From a geological perspective, losses of the AME peaked during the Jurassic, around 160 million years ago (Ma), followed by a decrease towards the early Cretaceous and another, smaller increase approximately 50 Ma. The larger peak coincides with the diversification of Synspermiata and the families Archoleptonetidae+Leptonetidae (Fig. 3b) and the smaller coincides with the diversification of Anapidae. The regain of the AME had a slight peak during the Cretaceous around 80-100 Ma, when the families Pholcidae, Plectreuridae, and Caponiidae diversified (Fig. 3b). Among the secondary eyes, the best-fitting models returned transition rates of 0.0005 (ALEs and PLEs) and 0.0007 (PMEs, Supplementary Table S1). Transitions between eye configurations occurred at low rates (<0.08) with the highest rates from the four-eyed morphologies (2SE and PE+1SE) to their respective two-eyed configurations (1SE and PE; rates 0.01 and 0.08, respectively) and from PE+2SE to PE+1SE (rate 0.02).

### Eye loss correlates with habitat

We detected significant correlations between eye loss and occupancy of low-light environments (LLEs), such as under rocks, in crevices, and in caves and subterranean habitats (Fig. 4, Supplementary Table S4). In most cases, the transition from eight to fewer eyes appears to have occurred simultaneously with, or after, lineage diversification within a given low-light habitat (Fig. 4b, Appendix 6). However, in Synspermiata, Archoleptonetidae+Leptonetidae, and more recently, Anapidae and Symphytognathidae, eye loss occurred long after occupancy of low-light microhabitats (Fig. 4b). Despite the overall correlation, the relationship between habitat and eye loss is complex, with possible interactions emerging between eye number, type, and configuration, phylogenetic affinity, and habitat type. Additionally, over 500 species (8.2% of those affected by eye loss) inhabited ‘normal’ light environments like vegetation.

The six-eyed state, predominantly represented by the loss of the AME (3SE), was mostly associated with epigean LLEs across Mygalomorphae, Synspermiata, Palpimanoidea, Araneoidea, and the RTA clade (Fig. 4a). However, this configuration also occurred in selected cave-affiliated groups and in Uloboridae, which includes species associated with vegetation (Fig. 4a). However, detailed habitat data was lacking for about half of affected species (Fig. 4c). Four-eyed species were mostly found in vegetation and represented by the Uloboridae (2SE configuration; Fig. 4a, c; Appendix 1, Fig. 3a, c, Fig. 4). Outside this family, most four-eyed species were associated to LLEs such as ground, leaf litter, and caves. Two-eyed spiders occurred in a range of habitats, although a significant proportion (44%) inhabited LLEs including leaf litter, insect nests, and caves (Fig. 4b, c). Eyeless spiders were entirely related to LLE occupancy (Hidden-Correlated model, AIC: 724.68, AICc: 725.99).

Over 90% of cases occurred in caves and other very dark habitats, such as subterranean microfissures. 1,264 of the 6,181 species affected by eye loss lacked sufficient data to infer a microhabitat.

## DISCUSSION

Our findings highlight the evolutionary lability of eye number in spiders, despite their typical visual system architecture being generally well-conserved through more than 300 million years of diversification. We found evidence for many independent origins of eye loss at varying phylogenetic depths, with a strong correlation to low-light environments, but detected complex interactions between the affected eye pairs, phylogenetic distribution, and specific habitats.

The modular nature of the spider visual system, i.e. having two eye types, one of which is present in triplicate, has important implications for the evolution and dynamics of eye loss. Our results indicate three overall trajectories: loss of the AME first; loss of the secondary eyes first; and the loss of all eyes near-simultaneously (Fig. 5).

### Loss of the AME

Our analyses indicate that the AME have been lost at least seven times independently in spiders and potentially regained three times within Synspermiata (Fig. 3a). This demonstrates far higher evolutionary lability in deep time than the secondary eyes, raising questions of both mechanism and causality.

The principal and secondary eyes develop from separate primordia that express different combinations of retinal determination genes (RDGs), as seen in the ocelli and compound eyes of insects (Homann 1971; Samadi et al. 2015; Schomburg et al. 2015; Baudouin-Gonzalez et al. 2022). The separation of these regulatory networks presumably allows independent, or semi-independent, evolutionary trajectories of loss between eye types. The loss or disruption of RDG networks specific to the principal eyes is a plausible mechanism for AME loss. However, RDG repertoires appear to be highly conserved across species with and without AME, and early expression of principal eye-specific RDGs persists even 160 Ma after their hypothetical loss in the synspermiatan *Segestria senoculata* (Baudouin-Gonzalez et al. 2022; Tong et al. 2025). Interestingly, pRNAi knockdowns of *sine oculis 1* (expressed in all eye primordia) predominantly affected AME development in *Parasteatoda tepidariorum* (Gainnet et al. 2020); although this could hint at greater vulnerability to disruption in the principal eyes, other factors including differential penetrance between eye types preclude this interpretation for now. Similarly, an apparent lineage-specific duplication of *orthodenticle*, expressed only in the developing principal eyes, was recently detected in Synspermiata, but no functional attempts have been made to connect this to AME instability in the group (Baudouin-Gonzalez et al. 2025).

Eye loss generally occurs under low-light conditions, when the energetic cost of eyes is no longer justified (Sumner-Rooney 2018; Porter 2025). Typical habitats include caves, the deep sea, and other environments where little light is available. We found a strong overall correlation between eye loss and occupancy of LLEs, but these are not overwhelmingly cave-living species; most spiders that have lost only their AME live in epigean environments. This suggests that the AME are readily lost under dim light, and not only in total darkness. They generally have greater acuity than the secondary eyes, and multiple wavelength sensitivities (Morehouse 2020), both of which confer the greatest benefit under higher light intensities. In the context of dim-light LLEs, visual needs may be more readily fulfilled by the secondary eyes, which are more often monochromatic and equipped with a reflective tapetum.

### Loss of the secondary eyes

The loss of secondary eye pairs was rare and exclusively occurred within genera or families. The retention of the AME alongside the loss of SEs was only reported in nine families.

The loss of selected pairs of secondary eyes could equally be explained by a failure of the eye primordium to split during the embryonic development, or by the degradation of individual eye pairs after splitting. The former hypothesis was tentatively proposed (Baudouin-Gonzalez et al. 2025) based on an unpublished observation that a species of *Tetrablemma* had multiple retinas beneath a single lens. We observed this phenomenon in another tetrablemmid, *Matta teteia*, and in the sole specimen of the diplurid *Harmonicon cerberus* (Appendix 2b). Other reports of eye fusion in individual specimens were considered teratological. The loss of eyes after primordium splitting is implied where the secondary eyes are reduced in size or visibly degraded. Two sister species in the palpimanid genus *Hybosidella* support this hypothesis: one has vestigial secondary eyes while the other has lost them completely (Wood et al. 2024). Additional examples appear in the genus *Caponina*, where the PMEs are lost, the ALEs are reduced, and the PLEs may be either lost or vestigial (Galan-Sanchez & Sumner-Rooney, unpublished data, Appendix 2h, i), and in multiple zodariids with reduced PMEs (Shafaie et al. 2025). The mechanism for losing all three secondary eye pairs has never been explicitly examined, but, as for the principal eyes, disruption or failure to initiate of the relevant RDGNs could cause the extinction of the secondary eye primordia.

Ecological correlates of secondary eye loss are unclear; we recovered support for the loss of the PLEs in low-light environments, and equivocal support for correlated and independent models of PME loss in LLEs, but were surprised that the relationships between light environment and loss were not stronger. Our ancestral state reconstruction showed that all losses of the secondary eyes are recent, occurring at the tips of the tree except for the PMEs in *Anapistula*. However, these analyses are limited to the resolution of the phylogeny (544 species), which may underrepresent secondary eye losses and their correlates. The availability of ecological data may also mask underlying patterns; for example, the loss of secondary eyes and retention of the AME predominantly occurred in Caponiidae and Palpimanidae, two distantly related groups with relatively little known about their ecology, behaviour, or natural history. Although their representation in the phylogeny was not sufficient to resolve it, the widespread loss of secondary eyes in both families suggests that they harbour ancient losses not captured by our analysis.

### Complete loss of eyes

Eyelessness was the second most common eye-loss phenotype in our study after the loss of the AME, i.e. 3SE. In stark contrast to principal eye losses, eyelessness is not associated with any high-level clades, occurring in (at least) 30 families across spider phylogeny (Fig. 2). Total eye loss occurred mainly at the genus and species level and was therefore not well captured by phylogenetic analyses. However, eyelessness was strictly associated with LLEs in the wider dataset (Fig. 4c), and over 90% of affected species were cavernicolous. Most caves are geologically recent, limiting the history of cavernicolous species to a few million years at most (Culver and Pipan 2019). The evolution of total eye loss therefore occurs rapidly, likely due to a combination of extreme selection pressures and genetic isolation. A recent preprint (Tong et al. 2025) found that, in two distantly related instances, the complete loss of eyes was associated with genome-wide intensified selection, while the loss of individual eye pairs coincided with relaxed selection in selected developmental genes. They detected similar selection dynamics in five specific genes across both instances of total eye loss, which they suggest indicates parallel evolution. While these findings provide a crucial first insight to the genomics of eye loss, the shift to cave living has enormous implications for many other aspects of physiology and morphology that may be reflected in these genomic changes. Finally, despite these strong drivers, eye loss is still not universal or immediate; remaining intact or vestigial eye pairs reported in current cave species may yet disappear completely in future generations.

### Distribution of eye losses

Although eye loss has occurred repeatedly across the major spider clades, losses are not uniformly distributed across the phylogeny, nor through geological time.

Several evolutionary hotspots are apparent within the phylogeny. Our ancestral state reconstruction highlights the early-diverging clade Synspermiata, which exhibits the loss and - surprisingly - multiple subsequent regains of the AME. Our broader survey captures enormous additional diversity in eye number not resolved by our phylogenetic analysis; four of the nine families exhibiting at least four different eye configurations are synspermiatan. In particular, Pholcidae and Caponiidae showed extraordinary diversity, including eight-, six-, four-, two-eyed, and eyeless species (Fig. 2, Appendix 1). However, these represent different evolutionary trajectories, as caponiids retain their AME, which are commonly absent in pholcids. This begs the question of whether, and how, eye loss (and regain) is adaptive in these groups. Factors such as habitat selection, activity patterns, prey capture, sexual selection, and predation pressures may all contribute to eye number, but the lack of reliable data on these points precludes formal analysis.

Our analyses identified multiple deep, ancient AME losses that largely corresponded to low-light environments (Fig. 3a, Fig. 4b). Interestingly, however, these did not occur uniformly through time, with peaks around 160 Ma and 50 Ma (Fig. 4b). The peak of AME losses coincides with a period of diversification and ecological success of several lineages during the Mesozoic, most notably Synspermiata, which represents 21% of fossil taxa at that time, compared to 13% today (Wunderlich 2008; Magalhaes et al. 2020). However, the shift in relative species richness between Synspermiata and entelegyne spiders from the Mesozoic to Cenozoic eras exemplifies a faunal turnover, during which AME loss also occurred independently in additional groups such as araneoid and RTA-clade families.

In contrast, more recent losses often occurred within cave-dwelling lineages and involved eyes of both types. Although we are unable to analyse correlations or evolutionary rates for these events, our wider survey provides valuable insights to how they are distributed across different eye configurations and taxonomic levels. For instance, the 2SE, 1SE, and eyeless configurations are widespread throughout the order and appear to constitute the dominant phenotype in specific genera (e.g., *Troglodiplura*, *Cicurina, Miagrammopes*).

Other unique cases of recent eye loss are characterised by the retention of the AME accompanied by the loss of one or more secondary eye pairs (Fig. 5, Appendix 1). Although this tends to occur at genus or species level, it can occur at higher taxonomic ranks; in Palpimanidae, for example, these losses are subfamily-specific (Wood et al. 2024), and within Caponiidae the loss of two secondary eye pairs coincides with the diversification of the subfamily Nopinae (Galan-Sanchez and Alvarez-Padilla 2022). Interestingly, the loss of secondary eyes with retained AME seems to have evolved since the mid-Cretaceous or later in all groups with this eye configuration (Adrian-Serrano et al. 2024; Wood et al. 2024; Gajski et al. 2025), but the functional significance and ecological correlates of these reductions remain unclear.

### Intraspecific variation

Sexual dimorphism is widespread and sometimes extreme in spiders. The only case we found of variation in eye number was an oonopid species that was described from a six-eyed male (lacking AME, typical for the family), but the depicted female clearly also lacks the PLEs (with no other visible malformation); however, this was not discussed in the study (Ott et al. 2019) (Appendix 2e, f). The degree of eye degeneration was sexually dimorphic in three cave species (Appendix 2c, d): two leptonetids, in which males’ eyes were well-developed, while females’ were vestigial (Wang et al. 2017), and conversely, one cicurinid with small but well-formed eyes in females, but almost eyeless males (Shimojana and Ono 2017). We found evidence of wider intraspecific variation on eye number and reduction in multiple cave-dwelling spiders, with some of them having colonized other environments (Supplementary Table S4). Precise accounts of intraspecific variation are rarely included in taxonomic descriptions, or species are described from short series of specimens. Even taxonomic misidentifications can be a confounding factor. Therefore, the observed patterns should be approached with caution.

### Impacts of eye loss

These data offer a starting point to explore the consequences of eye loss in spiders, which provide the opportunity to examine the impacts of eye loss on the remainder of the visual system. So far, just one study considered the impact of eye loss on the remaining eyes, in the four-eyed uloborid *Miagrammopes*. The loss of the anterior eyes is accompanied by the enlargement of the remaining retinae, rotation of the eye axes, and elevation of the PMEs and PLEs, allowing the animal to retain its dorsal and ventral FOV coverage despite losing FOV overlap between eyes (Opell and Cushing 1986; Opell and Ware 1987). Similar investigations in other groups exhibiting loss, especially those without a clear association to LLEs or another ecological explanation, will help us better understand the function of the spider visual system and its role in their ecological success.

### Beyond eye number

While we identify several interesting patterns in the evolution of eye number, there are other relevant aspects of visual system architecture this does not capture. One is eye size; eyes may reduce in size prior to loss, while the loss of selected eye pairs may be related to changes in the size of the remaining pairs. Recent studies show that eye pairs contributing to behaviours such as hunting tend to be enlarged and are often accompanied (possibly facilitated) by the reduction of other eye pairs (Chong et al. 2024). A possible extension of this trend could be the eventual loss of one or more eye pairs in parallel with increased investment in the remaining eyes. For example, a myrmecophagous clade of zodariid spiders have larger AME relative to body size (Gajski et al. 2025), which is usually accompanied by the reduction or loss of a secondary eye pair (PMEs). The size reduction of PME in salticids could also fit this explanation. Conversely, in cases trending towards total eye loss in very dark habitats, where loss is driven by energetic economy rather than reinvestment, the remaining eye pairs may also shrink.

### The evolution of eye number and losses

Spiders are just one example of visual systems with variable eye numbers. The ecological correlates of eye number variation have only previously been examined in large, distributed visual systems that have an indeterminate number of eyes (Audino et al. 2022); although these are not directly comparable to deviations from the relatively conserved eye number in spiders, correlations were still apparent between LLE occupancy and fewer eyes. Among arthropods, useful parallels might be drawn with the median ocelli of insects, or the variable number of lateral eyes in scorpions or myriapods. Insects may have three (ancestrally), two, or no ocelli, and although they are traditionally associated with flight (Ribi and Zeil 2018), losses of one or all ocelli appear in many insect groups, both flighted and non-flighted. To date, patterns of ocellus number have not been formally analysed, but this is a potentially fruitful area for future work. Within arachnids, the number of median eyes (the AME in spiders) is very stable, but the lateral eyes can vary substantially between and within orders. In particular, living scorpions have up to five pairs of lateral eyes, although attempts to homologise these suggest there are six major and four accessory ocelli found across the group (Loria and Prendini 2014). The number and configuration of the lateral eyes vary substantially between families, as well as within families and genera (Loria and Prendini 2014); this variation might be better captured by phylogenetic analysis and scorpions could provide an excellent model for examining lateral eye loss.

Our results reveal central evolutionary patterns on the mode and tempo of eye loss: a highly conserved conformation of eight eyes despite variability in eye number; a common loss of the principal eye system, which tends to occur at deeper nodes with significant ecological correlates in specific clades, and the recent, independent evolution of other eye loss reduction phenotypes. These results offer a foundation for future research into evolutionary patterns of eye number, reduction, and loss across different orders, their relationship with ecological and behavioural drivers, and their transformations through time.

## SUPPLEMENTARY MATERIAL

Data are available in the online version of this article [URL].

## ACKNOWLEDGMENTS

F.A.R.Q. thanks Jeremy A. Miller for his continuous support and for the constructive discussions on the early stages of this study. M.A.G.S. and L.S.R. are grateful to Russell Garwood and Jason Dunlop for their valuable feedback and enriching discussions on arachnid character evolution and spiders fossil record.

## DATA AVAILABILITY

The datasets generated and/or analysed during the current study are available in the Dryad Digital Repository xxx. All code used to analyse and present data is available via Zenodo: xxx

## AUTHOR CONTRIBUTIONS

M.A.G.S. developed the study concept, collected and analysed data, prepared figures, and wrote the initial draft of the manuscript. F.A.R.Q. developed the study concept, collected and analysed data, prepared figures, and contributed to writing. L.S.R. conceived the study, advised data analysis and visualisation, prepared figures, and contributed to writing.

## APPENDICES

**Appendix 1.**
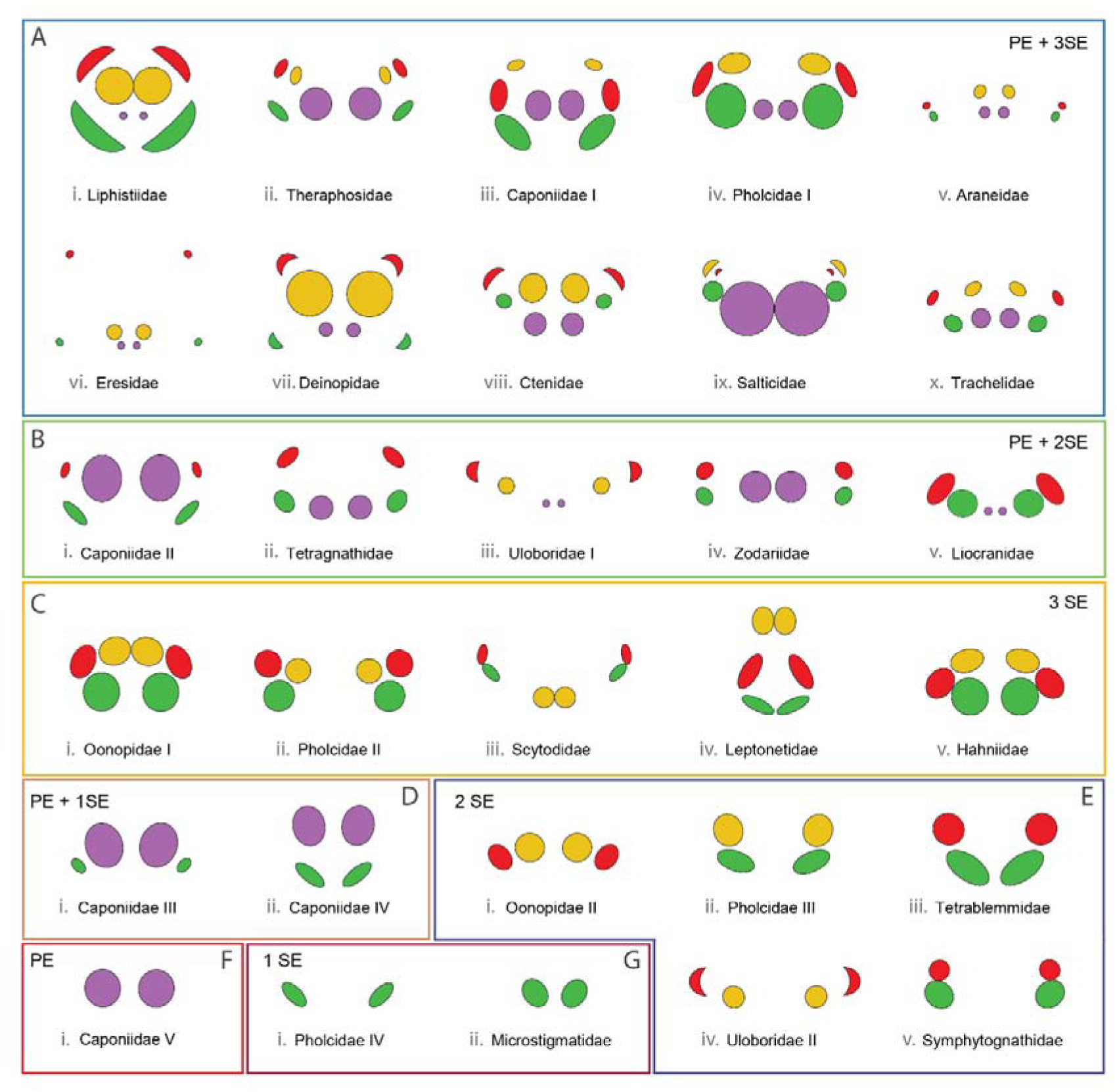
Diversity of the visual system in spiders. The stereotypical eight-eye morphology includes one principal eye pair (AME, purple) and three secondary eye pairs (ALEs, green, PLEs, red, and PMEs, yellow), which may be arranged in a variety of configurations with specific numbers and eye pairs. Colour frames exemplify the configurations analysed in this study. Note that some families exhibit multiple configurations (e.g., Caponiidae I-V or Pholcidae I-IV). Abbreviations: PE; principal eyes, SE; secondary eyes. After Morehouse et al. 2017

**Appendix 2.**
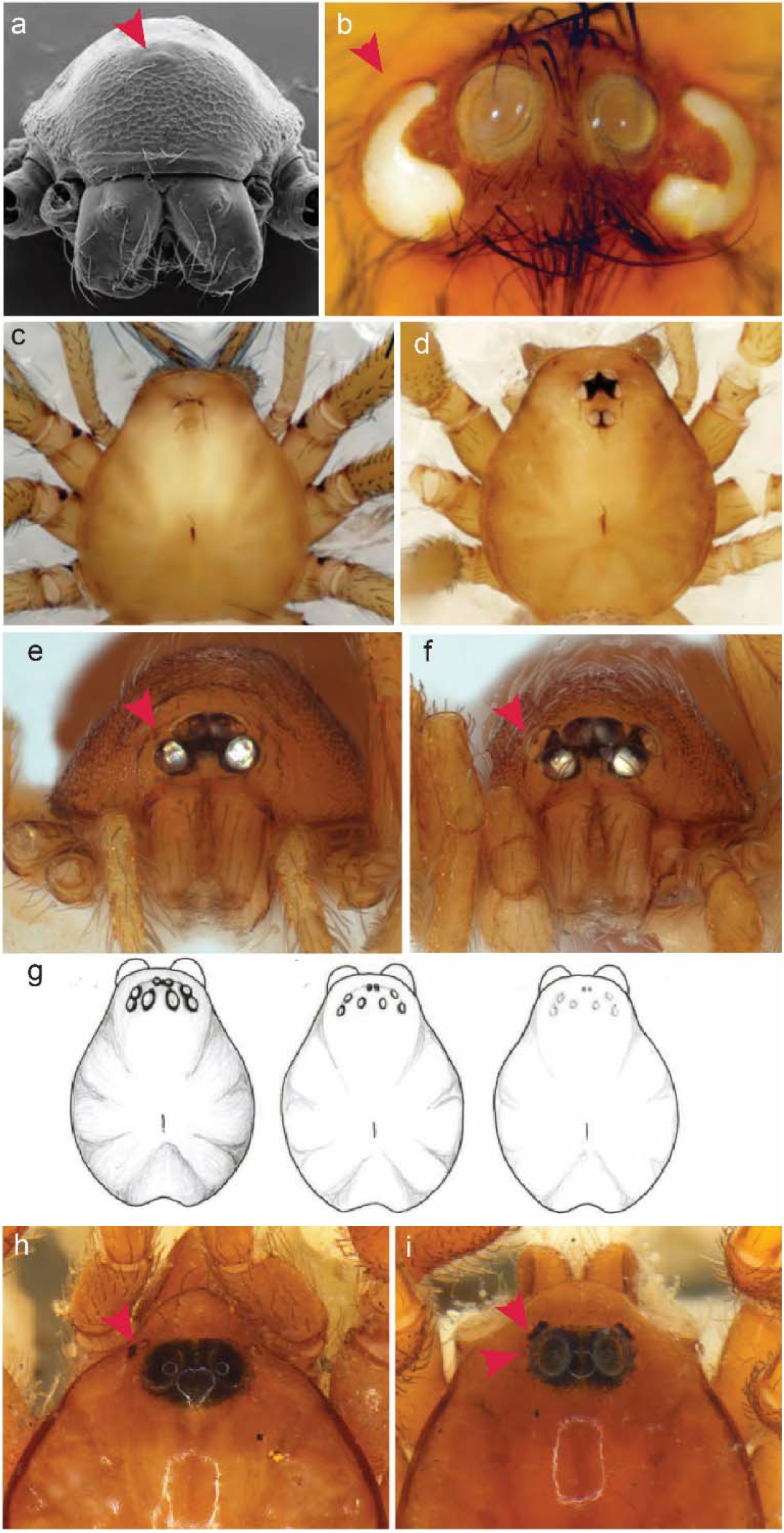
Eye fusion, eye reduction, sexual dimorphism, and intraspecific variation. a) The “one-eyed” tetrablemmid *Matta teteia* Brescovit & Cizauskas, 2019. b) The diplurid *Harmonicon cerberus* Pedroso & Baptista, 2014 whose SE have fused to form one elongated eye — albeit this species is only known from the holotype. c), d) The leptonetid *Leptonetela chenjia* Wang & Li, 2017, in Wang, Xu & Li, 2017, female paratype (left, with extremely reduced eyes) vs male holotype (right, with well-developed eyes). e), f) Sexual dimorphism in eye number in the oonopid species *Cinetomorpha campana* Ott & Harvey 2019, female paratype (left, with absent PLE) vs male holotype (right, with well-developed PLE). g) Variation in visual system development between different populations of *Troglohyphantes cantabricus* Simon, 1911 (from Fernández-Pérez et al. 2014). h), i) Degradation of secondary eyes in *Caponina* spp. h) The PME and PLE are absent and the ALE are highly reduced. i) The PME are absent and the ALE and PLE are reduced and highly reduced, respectively.

**Appendix 3.**
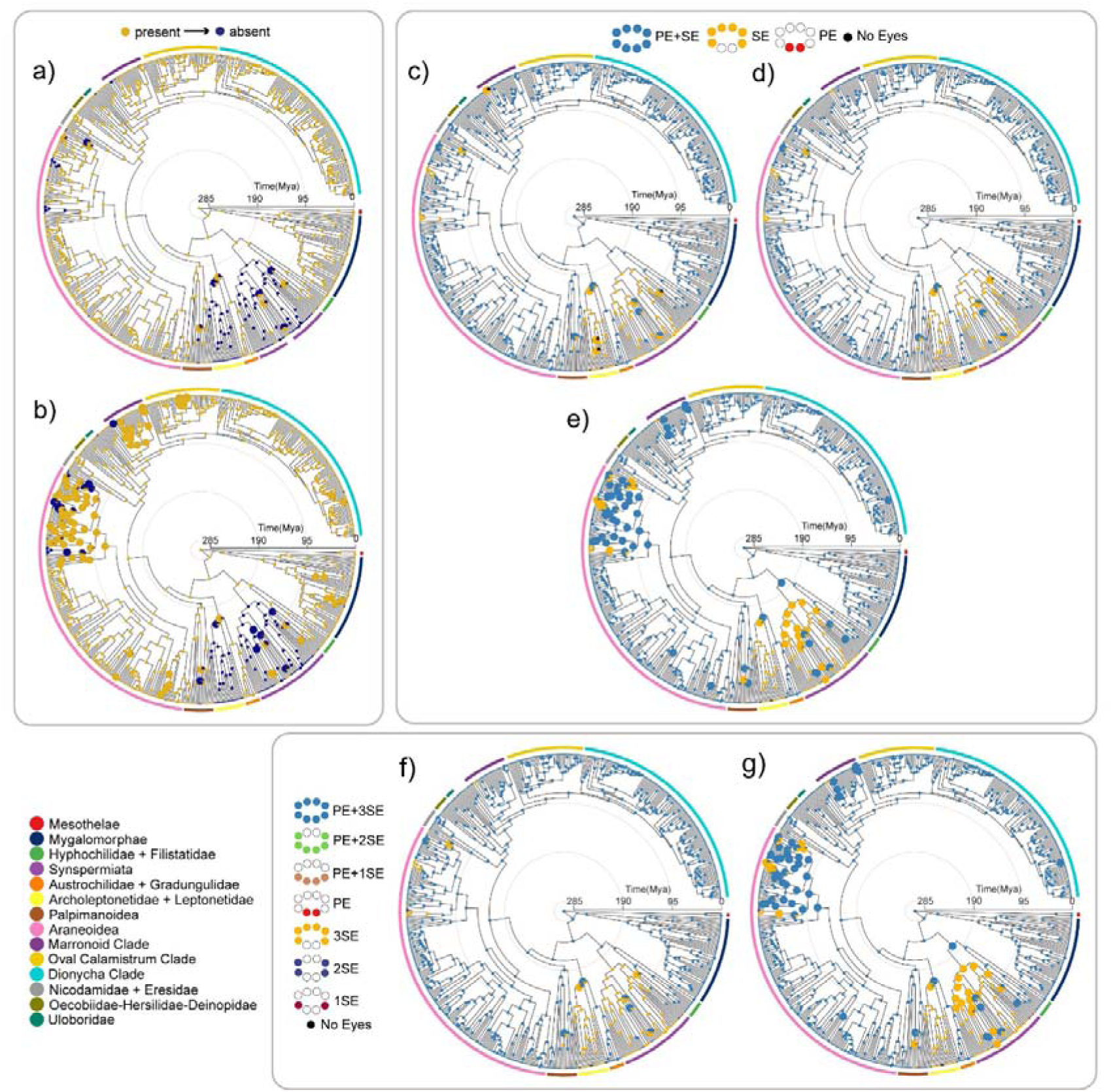
Marginal ancestral state reconstructions for eye loss under different character coding schemes. a), b) Ancestral state estimation of the anterior median eyes (AME) loss obtained under a) the 1-rate category (no hidden states) *Reversal ER* model and b) the 2 rate categories (one hidden state) *R1 No reversal - R2 Reversal ARD* model. c), d), e) Ancestral state estimation of eye loss analysing transitions from the presence of both visual systems (PE+3SE) to the absence of one of them (i.e., 3SE or PE) or both altogether (no eyes); ancestral state estimations obtained under c) the 1-rate category *Reversal SYM* model (no restrictions on matrix), d) the 1-rate category *Reversal ER* model (only PE+3SE -> 3SE and 3SE -> PE+3SE transitions allowed) and e) the 2-rate categories *Reversal ER - Reversal ARD* model. Transitions to PE and eyeless configurations have occurred relatively recently and appear only at the terminal tips in the phylogeny. f), g) Ancestral state estimation of eye loss analysing eight distinct configurations under f) the 1-rate category *Reversal ER* model and g) the 2-rate categories *Reversal ER - Reversal ARD* model. In both models only the PE+3SE -> 3SE and 3SE -> PE+3SE transitions are allowed. Note that only the 3SE configuration has evolved multiple times at deeper nodes in the phylogeny. The 2SE configuration has evolved at the *Anapistula* (Symphytognathidae) node. Other configurations are more recent although some may have evolved deeper in the diversification of some families (see discussion). Smaller charts at the nodes indicate probabilities > 90% for a given state. Abbreviations: PE; principal eyes, SE; secondary eyes.

**Appendix 4.**
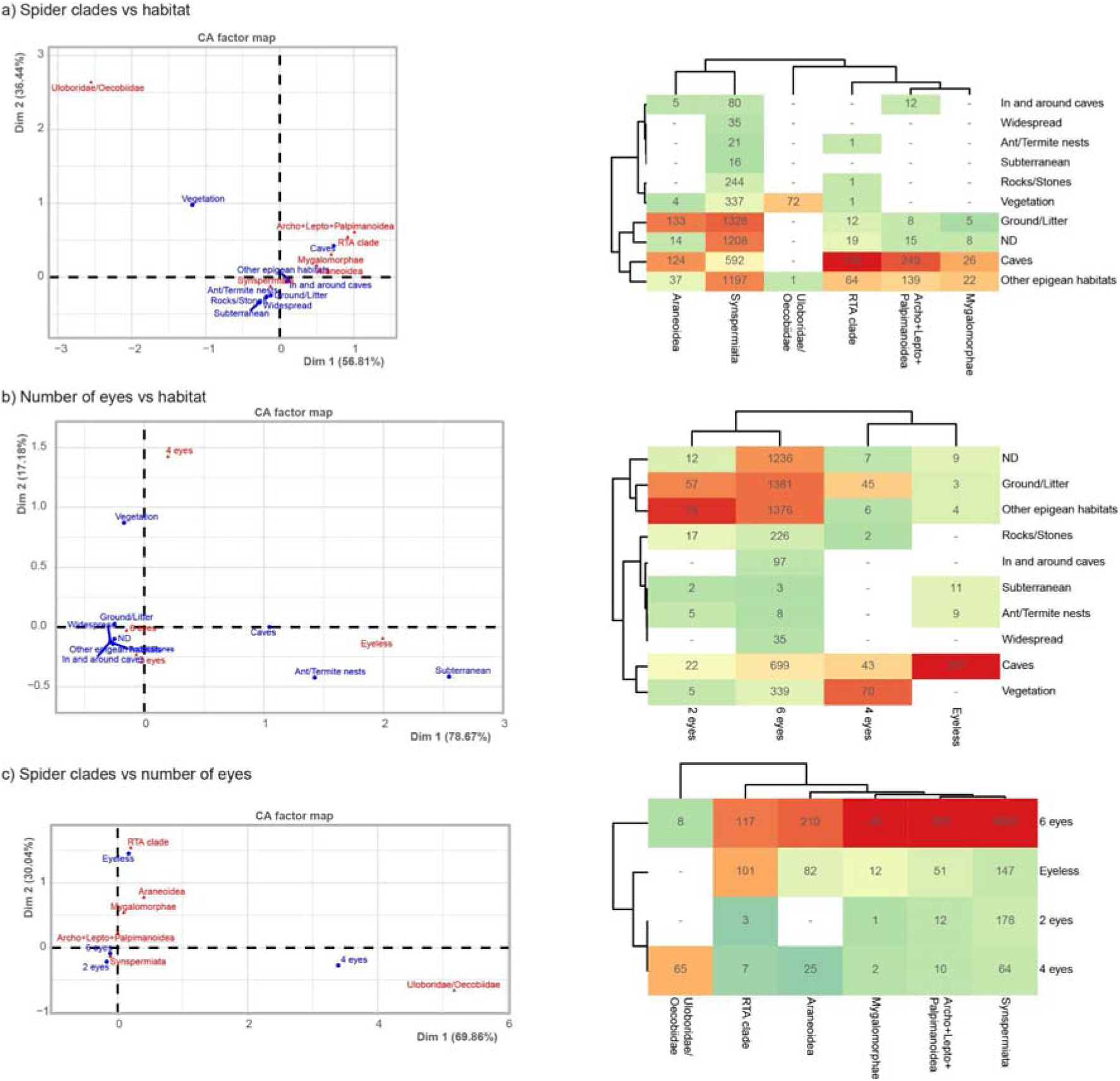
Relationships between eye loss and habitat occupancy across spider clades. a) Spider clade vs. habitat: Correspondence Analysis (CA) factor map (right) and heatmap (left). These visualizations demonstrate a strong affinity between specific clades and their habitats. For instance, Araneoidea, Mygalomorphae, the RTA clade, and Archo+Lepto+Palpimanoidea exhibit a higher specialization for cave environments than other clades, while Uloboridae/Oecobiidae is highly isolated and primarily associated with vegetation. b) Eye number vs. habitat: CA factor map (right) and heatmap (left) show a clear separation between photic and aphotic habitats. Eyelessness is strongly associated with life in caves, ant/termite nests, and subterranean habitats, while four-eyed morphologies are predominantly linked to vegetation. c) Spider clade vs. eye number: CA factor map (right) and heatmap (left) reveal a strong phylogenetic signal for eye morphology across spider clades. Eyeless species are more prevalent in Araneoidea, Mygalomorphae, and the RTA clade, while four-eyed morphologies are dominated by Uloboridae.

**Appendix 5.**
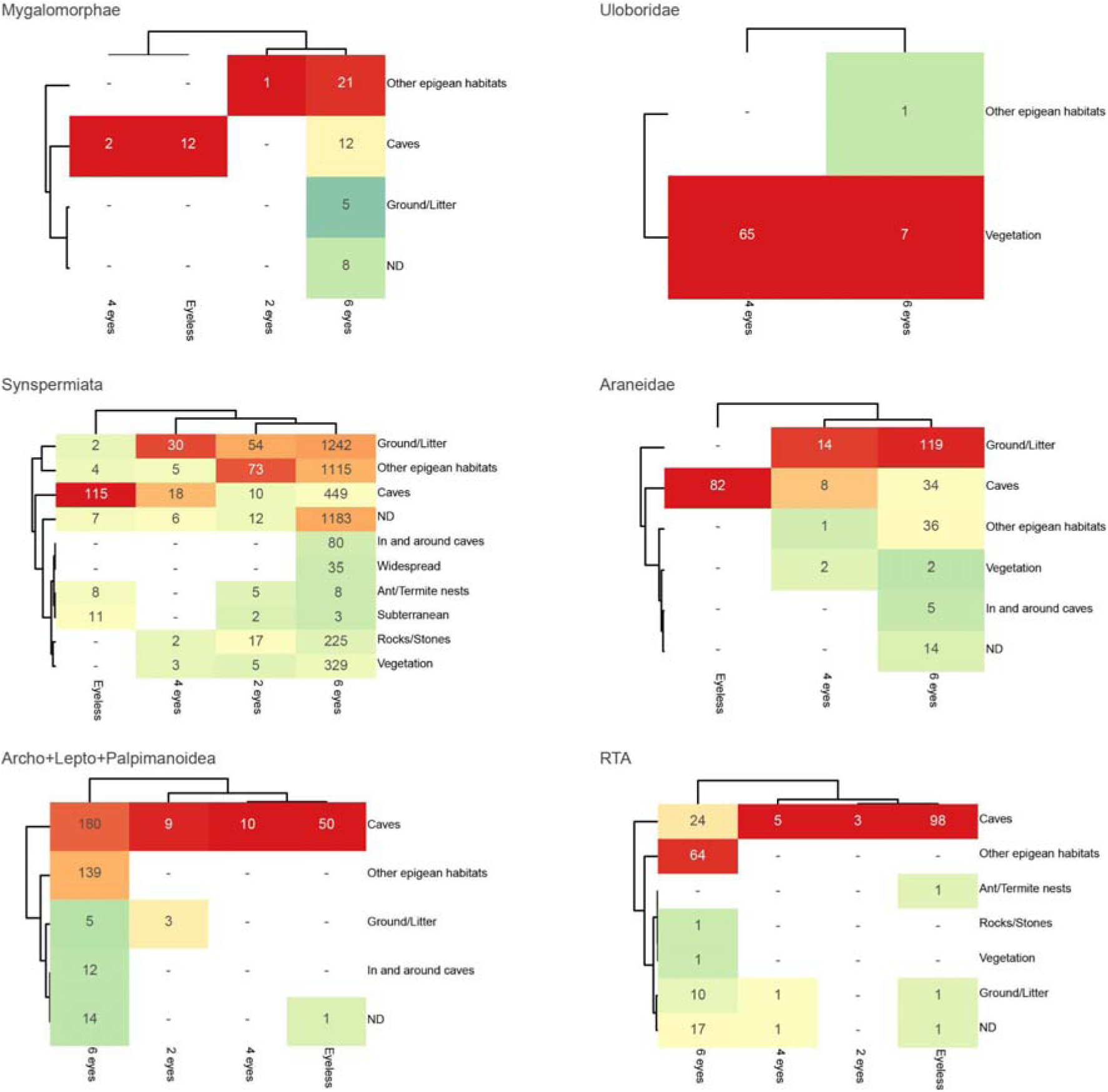
Relationships between eye number and habitat, detailed by spider clade. Note the uneven distribution of eye morphologies across all clades. In Mygalomorphae, the RTA clade, and the Archo+Lepto+Palpimanoidea, the majority of eye losses are associated with cave-dwelling. Uloboridae is dominated by four-eyed species associated with vegetation, while Synspermiata shows a significantly higher incidence of six-eyed species with no clear habitat correlation. Overall, the frequency of eye-reduced species in Synspermiata is notably higher than in any other clade.

**Appendix 6.**
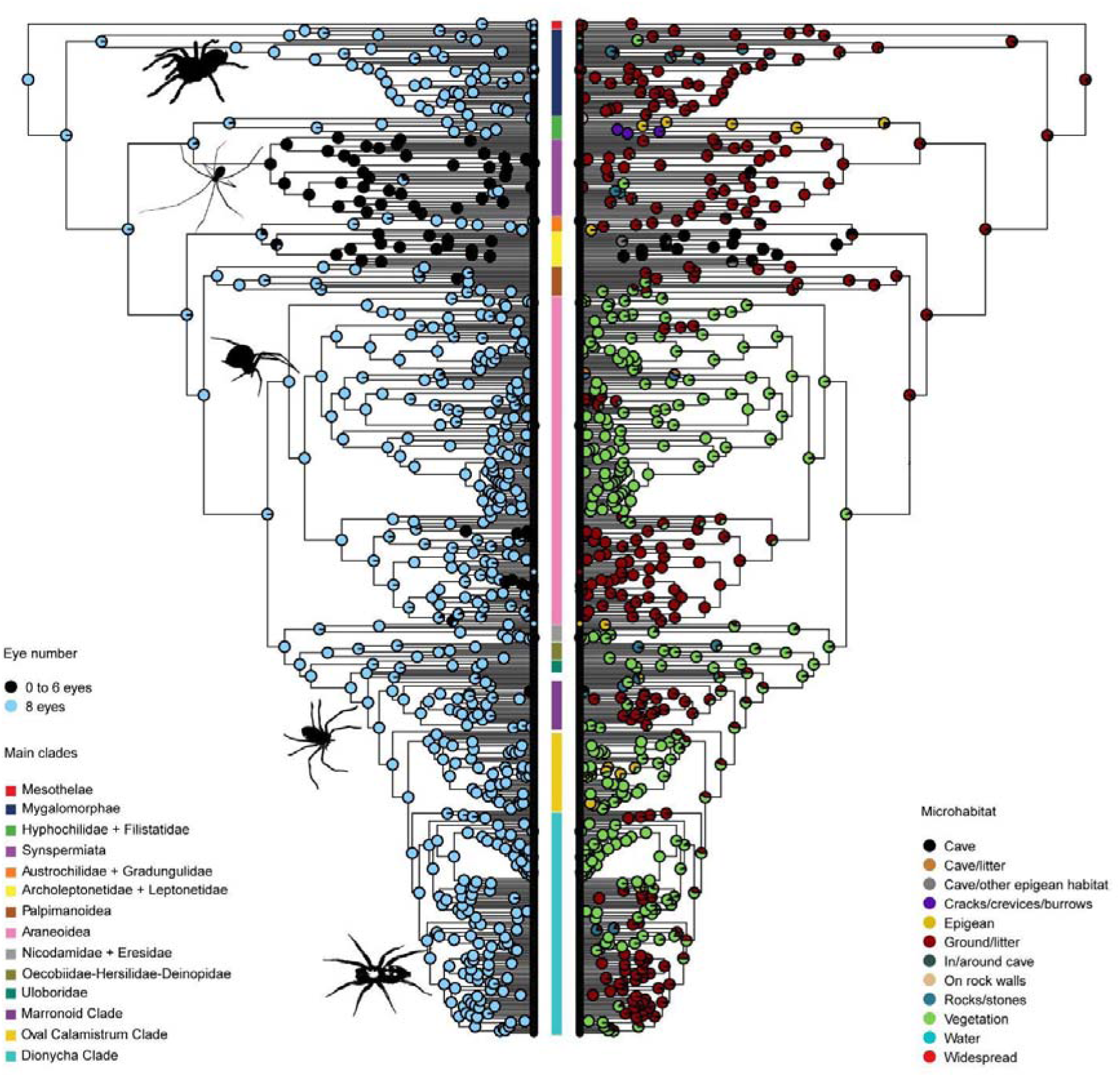
Maximum-likelihood phylogeny of spiders derived from a 25% occupancy dataset of the ultraconserved elements (Kulkarni et al. 2023), showing the ancestral state reconstruction of eye number (left tree) and microhabitats (right tree). Posterior probabilities at nodes of eye number were obtained under the best fitting ER model (AIC: 330.8790). Posterior probabilities at nodes of the ancestral microhabitats estimations were obtained under the best fitting SYM model (AIC:1310.0503). The inferred ancestral microhabitats for spiders include low light environments such as ground and litter with a shift to vegetation during the diversification of entelegynae spiders. Note, however, that within this clade, these environments were recovered as the ancestral habitats for the symphytognathoids, the Marronoid clade and families of the Dionycha clade.

